# Methylphenidate-induced reductions in choice impulsivity were associated with alterations in cortico-striatal beta oscillations

**DOI:** 10.1101/2025.09.18.675927

**Authors:** Jason Nan, Alyssa Terry, Noelle Lee, Morteza Salimi, Reyana Menon, Samuel A. Barnes, Dhakshin S. Ramanathan, Miranda F. Koloski

**Affiliations:** Mental Health, VA San Diego Medical Center, San Diego, CA, USA; Center of Excellence for Stress and Mental Health, San Diego, CA, USA; Department of Psychiatry, University of California-San Diego, San Diego, CA, USA

**Keywords:** temporal discounting, impulsivity, methylphenidate, cortico-striatal, beta oscillations

## Abstract

Impulsive choice- preferring small, immediate rewards over larger, delayed reward- is a hallmark of many psychiatric disorders. Methylphenidate, a dopamine and noradrenaline transporter inhibitor, reduces impulsivity, but the neural mechanisms underlying this effect remain unclear. Identifying a neurophysiological signature of methylphenidate’s action could be used clinically to predict treatment response and guide drug development. To this end, we recorded local field potentials (LFP) from 32 electrodes spanning cortico-striatal circuits in rats performing a temporal discounting task, under saline or methylphenidate treatment conditions. Methylphenidate decreased beta frequency (15-30 Hz) power and connectivity for small, immediate rewards while simultaneously increasing beta power for large, delayed rewards. Reduced connectivity in cortico-striatal networks during small rewards predicted increased preference for delayed rewards. These findings suggest that methylphenidate influences discounting behavior by modulating reward-evoked beta oscillations in cortico-striatal networks. Thus, beta oscillations may serve as a translational biomarker of treatment responsivity to monoaminergic drugs, providing a circuit-level assay to measure and predict changes in impulsive behavior.

## Introduction

Impulsive decision-making favors immediate actions over delayed rewards and is a core feature of psychiatric conditions including ADHD and substance use disorders (Ainslie, 1974; Dalley et al., 2011; Hamilton et al., 2015; MacKillop et al., 2011). Temporal discounting tasks, used across humans and animals, provide a translational assay of impulsivity that is sensitive to pharmacological manipulations (Ainslie, 1974; Baarendse & Vanderschuren, 2012; Dalley et al., 2011; Rodriguez & Logue, 1988; Winstanley et al., 2006). In discounting tasks, an impulsive choice describes the inability to wait for a larger reward over a small, immediate reward (Chudasama, 2011; Mitchell, 2019; Roesch & Bryden, 2011; Van Den Bos & McClure, 2013; Winstanley et al., 2005, 2006).

Methylphenidate, a dopamine and noradrenaline transport inhibitor, is considered a first-line treatment for ADHD and reduces impulsivity in both human and animal models (Castrellon et al., 2021; De Wit et al., 2000; Evenden & Ryan, 1996; Floresco et al., 2008; Pearson et al., 2004; Perry et al., 2009; Slezak & Anderson, 2011). Specifically, we have found that methylphenidate, and not citalopram (a selective serotonin reuptake inhibitor), increases preference for large, delayed reward on a temporal discounting task (Koloski, Terry, et al., 2024). However, the response to methylphenidate is variable (Caprioli et al., 2015; Rajala et al., 2015; Westbrook et al., 2020), likely reflecting individual differences in baseline catecholamine signaling (Clatworthy et al., 2009; Van Den Bosch et al., 2022). Identifying neurophysiological markers of methylphenidate’s effects would provide a translational assay to predict outcomes and personalize treatment (Arns et al., 2018; Sari Gokten et al., 2019).

Convergent evidence implicates cortico-striatal circuits in cost-benefit decision-making functions like temporal discounting (De Water et al., 2017; Kable & Glimcher, 2007; McClure et al., 2004). Top-down regulation of the ventral striatum via prefrontal and orbitofrontal cortex is associated with improved behavioral control and tolerance for delayed reward (Basar et al., 2010; Cardinal et al., 2002; Haber & Knutson, 2010; Peters et al., 2016; Van Den Bos et al., 2014; Winstanley et al., 2005; De Water et al., 2017; McClure et al., 2004). Altered cortico-striatal connectivity is observed with ADHD and other impulse-control disorders (Hong et al., 2015; Pujara & Koenigs, 2014). While methylphenidate reliably increases extracellular striatal dopamine (Faraone, 2018; Volkow, 1997), its effect on network-level activity underlying impulsive choice remains unclear.

Neural oscillations provide a dynamic readout of large-scale network interactions. Beta oscillations in prefrontal and striatal networks track reward expectancy, subjective value, and choice selection across species (Cohen et al., 2007; HajiHosseini & Holroyd, 2015; Hoy et al., 2023; Marco-Pallarés et al., 2015; Patai et al., 2022; Torrecillos et al., 2018; Xiao et al., 2024; Zavala et al., 2015). On a temporal discounting task, we have previously shown that reward-related beta oscillations are sensitive to temporal delay (Koloski, Hulyalkar, et al., 2024). We hypothesize that methylphenidate modulates reward-evoked oscillations to bias choice away from immediate, impulsive rewards.

Here, we use large-scale local field potential recordings in rats performing a temporal discounting task to test if cortico-striatal oscillations provide a mechanistic and translational marker of methylphenidate’s effects on impulsive choice.

## Materials and Methods

### Ethics Statement

This research was conducted in strict compliance with the Guide for the Care and Use of Laboratory Animals of the National Institutes of Health. The following protocol was approved by the San Diego CA Medical Central Institutional Animal Care and Use Committee (IACUC; Protocol Number A21-012).

### Animals

20 Long-Evans rats (10 males and 10 females) obtained from Charles River Laboratories were used in this study. We were powered for sex and initially tested for sex differences; however, sex was not a significant factor in behavioral results and therefore was not included in the final models (see Statistics). Rats were received at approximately one month old (150g) and began habituation/ pre-training two weeks after arrival. Rats were housed in pairs in standard cages (10 x 10.75 x19.5 inches) and kept on a 12h light/ dark cycle (6AM/6PM). Behavioral testing occurred during the light cycle. Animals were put on a water restriction schedule where they receive free access to water for 2 hours/ day to maintain motivation for water reward during operant behavior training. On non-training days, rats had free access to water and were weighed weekly to ensure that water restriction did not reduce food intake or lead to dehydration. Food was provided *ad libitum*.

### Experimental Design

Replicating our previous study (Koloski, Terry, et al., 2024), the experiment was designed to measure the dose-dependent effect of methylphenidate on temporal discounting behavior and its associated brain networks. Rats performed the temporal discounting task 5 days/ week. Rats received alternating intraperitoneal (i.p.) injections of methylphenidate (1.0, 5.0, or 10.0 mg/kg) and saline (control) injections. Between every drug/ saline pair, there were “off” days (no injections) to ensure adequate time between each drug session. Electrophysiology was recorded during every drug and saline session, but not during behavior only sessions. Drug was delivered in a pseudo Latin-squares design (rats experienced all doses in a randomized order). The temporal discounting task included two types of sessions: short delay (0.5s, 2s, 5s) and long delay (0.5s, 10s, 20s) sessions. Subjects first completed short delay sessions (two repeated measures of each dose) and then completed long delay sessions (two repeated measures of each dose). Two male rats did not reach pre-training requirements and were excluded from the study. 18 rats (8 male, 10 female) completed the experiment.

### Behavioral Training

#### Behavioral Apparatus

Behavior was shaped in custom-built operant boxes controlled using a Raspberry PI module with MATLAB software (previously described in Buscher et al., 2020). Each box housed two auditory tone generators, a house light for cues, 5 nose-ports each with an LED and IR sensor, and 5 peristaltic stepper motors to deliver water to each nose-port through clear tubing.

#### Pre-Training and Habituation

Animals began with habituation to the chamber and basic operant training to shape responses, under water restriction (access to water in task and 1 hr. following). In the first training program rats learned that a nose-port with an LED “on” signaled an available response port and that responding with a nose poke would trigger an immediate water reward (20 µL delivered at 10µL/s, 500ms after response). Animals advanced when they performed 100 trials in 60 minutes for three consecutive days. All animals advanced within a week of training. Next, rats trained on a version of the temporal discounting task where they could choose between a small (10µL delivered at 10µL/s) and large (30µL, delivered at 10µL/s) reward that were both delivered immediately (500ms after response) to shape the basic trial sequence and develop a preference for large reward. The drug study and final delay discounting task was initiated when rats developed a clear preference for large reward (≥70% large reward choices/session) and consistently performed 100 trials.

#### Delay Discounting Task

Generally, in a temporal discounting task subjects must choose between a low-value reward delivered immediately, or a higher-value reward delivered after a temporal delay. In our version of the task (described in further detail in Koloski, Hulyalkar, et al., 2024; Koloski, Terry, et al., 2024), subjects choose between a small reward (10µL) delivered immediately (0.5s after response) or a large reward (30µL) delivered after a variable delay (0.5s, 2s, 5s, 10s, 20s) (**Fig 1A**). The large reward delay always started at 0.5s delay (matching small reward) to establish large reward preference and incrementally increased after 45 trials in either a) short delay or b) long delay sessions (**Fig 1B**). In short delay sessions, large reward delay <45 trials =0.5s, 45-90 trials = 2s, >90 trials = 5s. In long delay sessions, large reward delay <45 trials = 0.5s, 45-90 trials = 10s, >90 trials =20s. Sessions were split this way to capture multiple delay lengths while still ensuring a substantial number of trials/ delay to record electrophysiology data (which requires averaging across a large number of trials to protect against individual trial variability) and limiting the number of injections given to each subject. Data was pooled across short and long delay sessions as we were interested in methylphenidate’s effect across all 5 delay conditions. A 5s inter-trial-interval was initiated after a response. Each session lasted for 60 min and trials were self-paced. On average, rats performed 130 trials per session. Sessions with <70 trials were not included.

**Figure 1.**
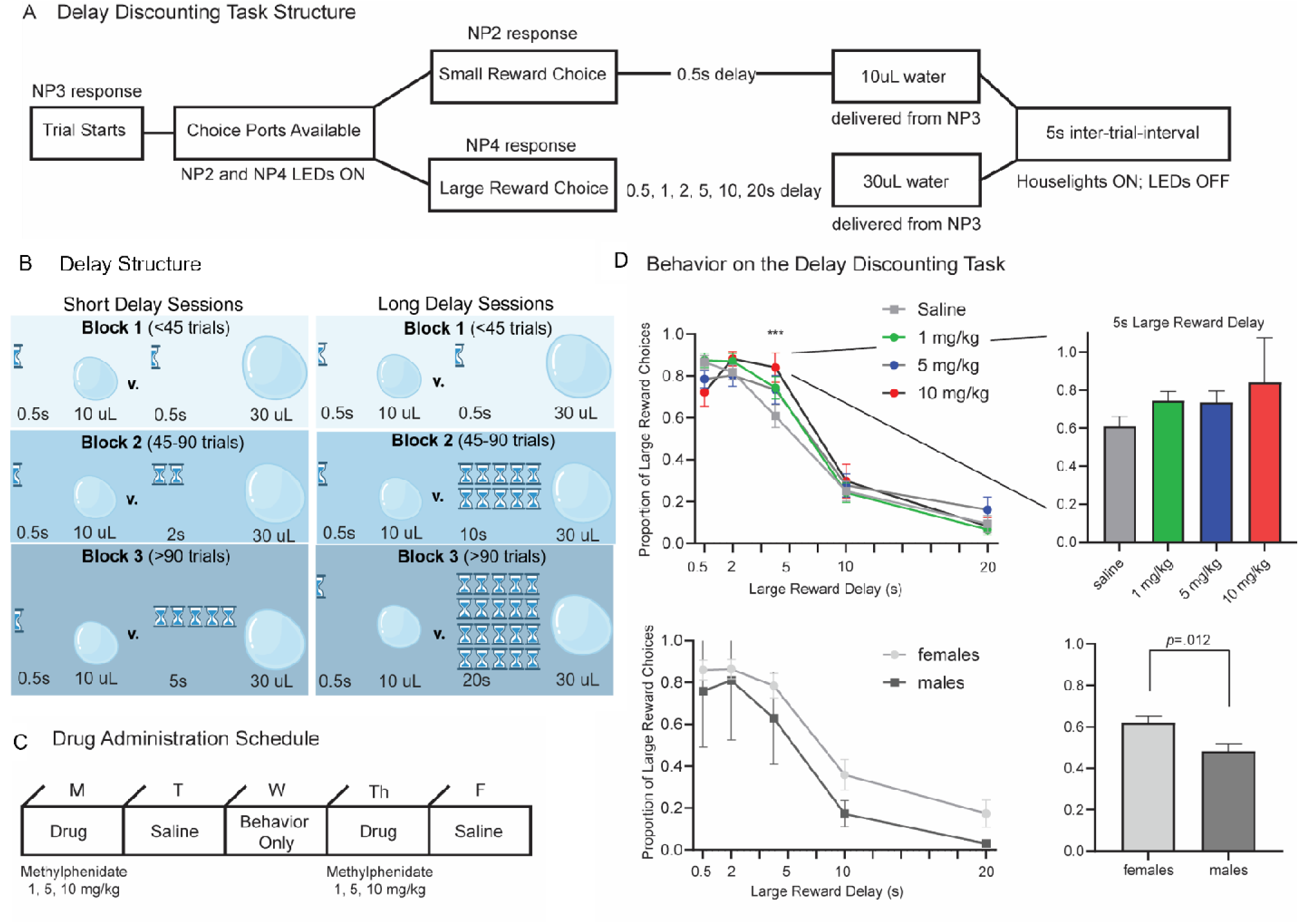
Task structure and behavior on the delay discounting Task. A) Schematic of the delay discounting task structure. Subjects chose between a small, immediate reward (10µL water delivered after 0.5s) or a larger reward (30µL water) delivered after a variable delay (0.5s -20s). The task was self-paced and lasted 60min. B) The large reward delay was variable and increased incrementally throughout the session. In short delay sessions, the large reward delay increased from 0.5s, 2s, 5s every 45 trials. In long delay sessions, the large reward delay increased from 0.5s, 10s, 20s every 45 trials. The timer represents the delay (half timer = 0.5s). Data was compiled across sessions for analysis. C) Methylphenidate (1, 5, 10 mg/kg, i.p.) or saline was administered according to a pseudo-latin squares design, to randomize drug dose order for each subject. Injections were administered 20 min before behavioral testing. Drug and saline days alternated, and a behavior-only day was interleaved to avoid carryover effects. D) Proportion of large reward choices across delay conditions (n=18 rats). Mean ± SEM are shown. At the shortest delay (0.5s), rats showed a strong preference for the large reward (86.6 ± 3.1%). As delays increased, choice of the large reward decreased (main effect of delay; F_(4,273.55)_ = 260.791, *p* <*.001*)- an effect that was mitigated by methylphenidate (dose x delay interaction; F_(12, 273.24)_ =2.90, *p<.001*). The effect of drug was most significant at the 5s delay (post-hoc one-way ANOVA; F_(3,60)_ =2.82, *p=.041*). Discounting behavior is also stratified by sex. Females (n=10) were less impulsive than males (n=8) (main effect of sex; F_(1, 16.01)_=7.98, *p=.012*), but no sex x drug interactions were observed (F_(12, 273.24)_ =1.01, *p=.442*) and therefore sex was not explored further.

#### Drug Administration

We used injections (i.p.) of methylphenidate (Sigma-Aldrich, St. Louis, MO, USA) at 3 doses (1, 5, 10 mg/kg) delivered in a pseudo-Latin squares design. These doses have been used in various studies to assess impulsive behavior while minimizing the risk of inducing stereotypical motor behaviors (Roffman & Raskin, 1997; Yang et al., 2006). As a control, saline injections were given the session prior to any drug session (**Fig 1C**).

Methylphenidate was mixed in 0.9% saline and injected at a final concentration of 1mL/kg. As a control, 1mL/kg 0.9% sterile saline was administered. IP injections were given 20 minutes before the start of a behavioral session to allow methylphenidate to reach its peak concentration (15-30 min),yet still have a duration (60-100 min) that would last throughout the session (Askenasy et al., 2007).

### Electrophysiology

#### Electrode Fabrication and Design

The fabrication and implantation procedures of our custom 32-CH local field potential (LFP) probes are described in detail (Francoeur et al., 2021). Briefly, we arrange electrodes into 8 cannula (“bundles” of 4 electrodes) to target 32 locations simultaneously (one electrode per region). 50µm tungsten wire (California Fine Wire, CA, USA) is arranged in “bundles” of 4 wires and glued in 30-gauge stainless steel metal cannula (Mcmaster-Carr, Elmhurst, IL, USA) cut 8-9mm long. Each electrode wire is cut to their unique D/V length. The 8 cannulas are positioned to target different AP/ML locations. **Table 1** provides the AP/ML/DV of each target region. The average impedance of our electrodes is 50 kOhms at 1 kHz.

**Table 1.** Electrode locations and reward-evoked power. Name, abbreviation and coordinates (AP,ML, and DV) are shown for each electrode based on Paxinos & Watson atlas, 2013. Normalized beta power during reward (0-1s from reward onset) is reported for each electrode on small and large (0.5s delay) trials. Significance (t-test from null) is denoted with asterisks (* *p*<.05, ** *p*<.01; \*\*\**p*<.001).

| Electrode Name | Abbreviation | AP | ML | DV | Small Reward | Large Reward (0.5s) |
| --- | --- | --- | --- | --- | --- | --- |
| Anterolateral motor cortex | M1 | 3.75 | 3.2 | 1.0 | 0.95 ± 1.31 *** | 0.74 ± 1.01 *** |
| Lateral frontal cortex | LFC | 3.75 | 3.2 | 3.6 | 0.37 ± 0.92 * | 0.24 ± 0.96 |
| Anterior insula | Ains | 3.75 | 3.2 | 4.8 | 0.45 ± 1.00 * | 0.44 ± 1.13 |
| Lateral orbitofrontal cortex | LOFC | 3.75 | 3.2 | 5.8 | 1.07 ± 1.33 *** | 0.80 ± 1.19 *** |
| Secondary motor cortex | M2 | 3.75 | 0.8 | 0.8 | 0.66 ± 1.16 ** | 0.53 ± 1.14 |
| Dorsal prelimbic cortex | A32D | 3.75 | 0.8 | 3.2 | 0.35 ± 0.69 ** | 0.23 ± 0.90 |
| Ventral prelimbic cortex | A32V | 3.75 | 0.8 | 4.8 | 0.44 ± 0.73 ** | 0.30 ± 1.12 |
| Ventral orbitofrontal cortex | vOFC | 3.75 | 0.8 | 5.8 | 1.08 ± 1.23 *** | 0.82 ± 1.64 *** |
| Motor cortex | M1 | 2.0 | 1.8 | 1.5 | 0.23 ± 0.85 | 0.04 ± 0.83 |
| Dorsomedial striatum | DMS | 2.0 | 1.8 | 4.5 | 0.61 ± 1.16 ** | 0.47 ± 1.59 ** |
| Ventromedial striatum | VMS | 2.0 | 1.8 | 5.7 | 0.73 ± 1.21 *** | 0.60 ± 1.41 ** |
| Nucleus accumbens core | NAcC | 2.0 | 1.8 | 6.9 | 0.69 ± 1.58 * | 0.59 ± 1.01 ** |
| Anterior cingulate cortex | A24b | 2.0 | 0.6 | 2.0 | 0.46 ± 0.93 ** | 0.44 ± 0.97 |
| Anterior cingulate cortex | A24a | 2.0 | 0.6 | 3.0 | 0.56 ± 0.93 *** | 0.44 ± 0.93 * |
| Anterior cingulate cortex | A33 | 2.0 | 0.6 | 3.5 | 0.29 ± 1.02 * | 0.21 ± 0.78 |
| Nucleus accumbens shell | NAcSh | 2.0 | 0.6 | 6.6 | 0.92 ± 1.26 *** | 0.74 ± 1.35 *** |
| Dorsolateral striatum | DLS | -2.5 | 4.9 | 5.1 | 1.20 ± 1.52 *** | 0.91 ± 1.76 *** |
| Dorsolateral striatum | DLS | -2.5 | 4.9 | 6.1 | 0.69 ± 0.81 *** | 0.57 ± 1.46 ** |
| Central amygdala | CEA | -2.5 | 4.9 | 7.1 | 0.75 ± 1.41 ** | 0.68 ± 2.08 |
| Basolateral amygdala | BLA | -2.5 | 4.9 | 8.1 | 1.14 ± 1.12 *** | 0.89 ± 1.62 *** |
| Cingulate cortex | A30c | -2.5 | 0.7 | 1.3 | -0.77 ± 0.97 *** | -1.03 ± 0.91 *** |
| Cingulate cortex | A29c | -2.5 | 0.7 | 2.3 | -0.79 ± 0.95 *** | -1.16 ± 0.63 *** |
| Mediodorsal thalamus | MD Thal | -2.5 | 0.7 | 4.7 | -0.57 ± 1.06 ** | -0.62 ± 1.22 ** |
| Centro-median thalamus | CE Thal | -2.5 | 0.7 | 5.7 | -0.80 ± 1.11 *** | -1.20 ± 0.87 *** |
| Posterior parietal cortex | PPC | -3.5 | 2.5 | 1.4 | -0.28 ± 1.05 | -0.38 ± 1.10 |
| Hippocampus | CA1 | -3.5 | 2.5 | 2.5 | -0.28 ± 1.15 | -0.38 ± 1.03 |
| Hippocampus | CA3 | -3.5 | 2.5 | 3.5 | -0.28 ± 1.27 | -0.40 ± 0.79 * |
| Subthalamic nucleus | STN | -3.5 | 2.5 | 8.0 | -0.19 ± 0.78 | -0.30 ± 1.20 * |
| Primary visual cortex | V1 | -6.0 | 3.5 | 1.0 | 0.07 ± 0.90 | -0.18 ± 1.31 |
| Primary visual cortex | V1 | -6.0 | 3.5 | 1.7 | -0.09 ± 0.99 | -0.30 ± 1.30 |
| Dorsal subiculum | DorSub | -6.0 | 3.5 | 2.8 | 0.26 ± 1.18 | 0.21 ± 1.04 |
| Dentate gyrus | DG | -6.0 | 3.5 | 3.7 | -0.34 ± 0.98 ** | -0.41 ± 1.26 * |

#### Surgery

Aseptic surgeries were performed under isoflurane (1-3%) anesthesia (SomnoSuite, Kent Scientific, CT, USA) in a stereotaxic frame with all instruments autoclaved prior to start. Core body temperature was maintained at 37°C ± 0.5°C using a temperature-controlled heating pad. Animals received a single dose of Atropine (0.05 mg/kg) to diminish respiratory secretions during surgery, a single dose of Dexamethasone (0.5 mg/kg) to decrease inflammation, and 1mL of 0.9% sodium chloride solution prior to surgery. The area of incision was cleaned with 70% ethanol and iodine solution. A local anesthetic, Lidocaine (max .2cc), was injected under the skin at the incision site while the animal was anesthetized but before surgery initiation. An incision was made to expose the skull, muscle and connective tissue cleared, and skull dried prior to drilling holes. During surgery, 8 holes were drilled in the skull (one for each cannula) at predetermined stereotactic locations (**Fig 2A**). Additional holes were drilled for a ground wire above cerebellum and anchor screws (5) around periphery.

**Figure 2.**
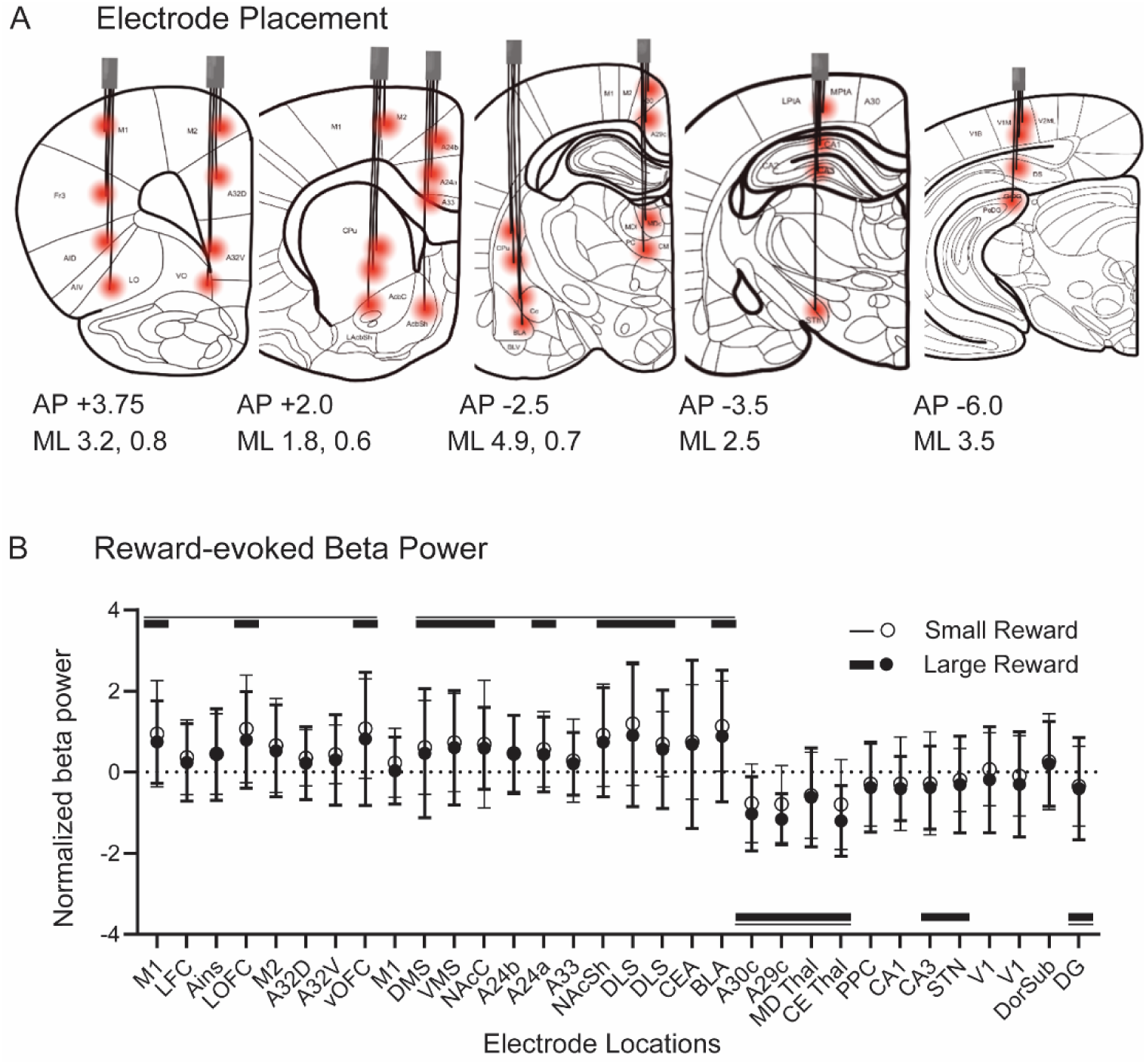
Electrodes of interest. A) Target locations of the 32 electrodes (red) displayed on representative coronal slices (modified from Paxinos & Watson, 2013). Four microwires were housed in a single cannula, each aimed at a unique AP/ML coordinate (**Table 1**). B) Reward-evoked beta power (15-30Hz, 0-1s from reward onset) plotted by electrode location in anatomical order (as shown in A). Mean ± SD normalized beta power is shown for small rewards (open circles) and large rewards (filled circles) when delays were equal (0.5s). Electrodes with normalized beta power significantly different from 0 (one-sample t-test, Bonferroni corrected) were classified as activated (above baseline) or suppressed (below baseline). Horizontal lines above or below indicate electrodes with significant activation/ suppression (p<.05) for small reward (thin line) or large reward (thick line). Only electrodes that were activated for both reward conditions were included in subsequent analyses (10 electrodes, **Table 1**).

Electrodes were slowly lowered to desired depth, pinned to the EIB board, and secured with Metabond (Parkell, NY, USA). The entire head stage apparatus was held to the skull and encased with dental cement (Stoelting, IL, USA). Once the head stage was secured, the skin was sutured closed, and rats were given meloxicam (1mg/kg) for pain management for 3 days following surgery. Rats recovered from surgery on a heating pad to control body temperature and received sulfamethoxazole and trimethoprim in their drinking water (60mg/kg per day for 8 days) to prevent infections.

#### Recording Design

LFP data was recorded using a 32-channel RHD headstage (Intan Technologies, CA, USA; Part C3324) coupled to a RHD USB interface board (Intantech, Part C3100) and SPI interface cable. We used plug-in GUI (Open Ephys) software for acquisition. Data was recorded at 1Khz, with a band-pass filter set at 0.3 to 999 Hz during acquisition. Physiology data was integrated with behavioral data using a lab-streaming-layer (LSL) protocol as described previously (Buscher et al., 2020).

#### Signal Processing

Local field potential (LFP) signals were preprocessed using a standardized pipeline to reduce noise and ensure data quality (see GitHub repository for full code: https://github.com/jasonnan2/Automated_LFP_Analysis). Each recording was segmented into trials aligned to behavioral events (-3 to +3s around reward onset). -1 to +1s from trial start was used for baseline normalization. Preprocessing included four steps:

1. **Noise removal:** Periods of broadband recording noise (high amplitude noise across all channels) were automatically detected and excluded for the entire time series.
2. **Channel rejection:** Electrodes with persistently abnormal activity (e.g., high noise or no signal from head stage/tether damage) were identified based on power spectra and removed. Any channel with over 30% frequency points > 3 median absolute deviations above activity of all channels was removed in an iterative manner until no channels are rejected.
3. **Trial rejection (low power):** Trials with abnormally low spectral power (average dB power < 0), reflecting poor signal, were discarded.
4. **Trial rejection (outliers):** Trials with unusually large voltage fluctuations (>3 SD) were excluded to remove artifacts from movement or recording instability.

After cleaning, signals were re-referenced within each electrode bundle (4 electrodes per cannula) to minimize the influence of volume conduction. On average, ∼94% of trials per session were retained (range: 72 to 99.7%), 29 channels per session were retained (range: 23 to 31), and ensuring adequate data for spectral analyses.

#### Power Analysis

Power analysis was done for each session with at least 10 trials of each delay condition (small reward, 0.5, 2, 5, 10, 20s delays) after trial rejection (above). The steps for power analysis are as follows, repeated for each subject, session (including all drug conditions), and delay condition:

1. LFP trial data is bandpass filtered for beta band (15-30Hz) through EEGLAB.
2. Power is calculated using pwelch function (hamming window with 50% overlap) and converted to dB.
3. Baseline correct each trial (-1s trial onset).
4. Calculate reward time window (+1s reward onset).
5. Calculate median across all trials.
6. Calculate median across all sessions.
7. Calculate average across all subjects.

#### Network Connectivity Analysis

To look at overall network connectivity metrics, we calculated weighted phase lag index (wPLI) using the fieldtrip package. For electrodes of interest (electrodes with significant beta activation during reward), we calculated their wPLI with the LOFC electrode (a cardinal region for reward processing where beta activation is the strongest). The same 8 procedural steps for calculating power (above) were used to calculate wPLI across 10 electrode pairs besides the baseline correction step which is not necessary for connectivity analysis.

### Statistical Analyses

#### Behavioral

Our primary behavioral measure was the proportion of large reward choices made at each delay. We examined trial count/ session as a secondary dependent measure. We used linear mixed models to account for missing sessions and variability between subjects (fixed factors [drug condition (saline, 1, 5, 10mg/kg), delay (0.5, 2, 5, 10, 20s), session], and random factors [subject, dose order]). Session was a repeated measure. To determine best fit of the model, we measured the Akaike information criteria (AIC) and Bayesian information criterion (BIC) of four commonly used covariance models (compound symmetry, scaled identity, AR(1), and unstructured (49.50). The scaled identity model, assuming repeated measures may be independent but with equal variance, provided the lowest AIC and BIC scores. A *p-value* less than 0.05 was considered significant. Significant results were followed with Bonferroni-corrected post-hoc comparisons (two-tailed). Both sexes were included and initially tested for sex differences; however, sex was not a significant factor in behavioral measures and therefore was not further analyzed for electrophysiological effects or included in the final models. Behavioral data was analyzed with IBM SPSS Statistics v.28 (New York, USA) and visualized in Graph Pad Prism v.9.

#### Electrophysiological

Statistical analyses on LFP power were similarly analyzed using linear mixed models. Analyses were performed separately for small, immediate rewards and large, delayed rewards. Electrodes with normalized beta power significantly greater than 0 (one sample t-test) in both small and large reward conditions (0.5s delay) were selected for further analyses. On small, immediate reward trials, beta power (dependent measure) was analyzed with a linear mixed model (fixed factors [drug condition (saline, 1, 5, 10mg/kg), electrode, session], and random factors [subject, dose order]). On large, delayed reward trials, beta power (dependent measure) was analyzed with a linear mixed model (fixed factors [drug condition (saline, low, medium, high), delay (0.5, 2, 5, 10, 20s), electrode, session], and random factors [subject, dose order]). A *p-value* less than 0.05 was considered significant. Significant results were followed with Bonferroni-corrected post-hoc comparisons. Similar linear mixed models were used to analyze weighted phase lag index (wPLI), a measure of network connectivity, as a dependent measure. wPLI was analyzed between LOFC and the other electrode sites of interest (10 connectivity pairs, corrected for multiple comparisons). Finally, to quantify the relationship between physiology and behavioral response, Pearson’s correlations were run between wPLI and proportion of large reward choice for each session and fit to a simple linear regression. Behavioral data was analyzed with IBM SPSS Statistics v.28 (New York, USA) and visualized in Graph Pad Prism v.9.

## Results

We used a large-scale recording technique to investigate neural signals contributing to the behavioral effect of methylphenidate on impulsivity. Previously, we have described beta oscillations in the cortico-striatal network that signal subject value on the temporal discounting task, decreasing their power and connectivity as the delay to reward increases (Koloski, Hulyalkar, et al., 2024). Here, we wanted to know if 1) there was an effect of methylphenidate on this putative beta reward signal and 2) if there was a physiological effect, was it reflecting behavioral changes in impulsivity?

### Behavioral Results

18 Long-Evans Rats (8 male; 10 female) performed a delay discounting task with fixed, but variable large reward delays while we simultaneously recorded brain-wide LFP signals (**Fig 1A**). Based on our previous work showing a dose-dependent effect of methylphenidate to increase large reward preference at long delays (Koloski, Terry, et al., 2024), we administered low (1 mg/kg), medium (5 mg/kg), and high (10 mg/kg) doses of methylphenidate during delay discounting (**Fig 1B, C**). First, we used a linear mixed model, to see how methylphenidate injections change impulsive choice at different delay lengths (0.5, 2, 5, 10, 20s). As expected, the proportion of large reward choices declined as the temporal delay to reward increased (main effect of delay; F_(4,273.55)_ = 260.791, *p* <*.001*) (**Fig 1D**). Under control conditions (saline injection), rats strongly preferred the large reward choice when delays were equal (0.5s delay (86.6+/- 3.1%) and discounted the large reward at increasing delays (20s delay (9.4 +/-16.5%). Methylphenidate increased the likelihood of choosing the large, delayed reward in a dose-dependent manner, particularly at the 5s delay (dose x delay interaction; F_(12, 273.24)_ =2.90, *p<.001*) (**Fig 1C**). Females were generally less impulsive than males (main effect of sex; F_(1,16.01)_=7.98, *p=.012*), but the drug’s effect did not differ by sex (*p=.442*), so data were pooled.

### Beta Frequency Activation for Reward

Next, we explored oscillatory activity that may explain the observed effect of drug. We employed a data- driven approach to identify brain activity susceptible to dopaminergic modulation via methylphenidate. We defined regions of interest based on reward-evoked beta power (15-30 Hz), which we previously found to reflect subjective value of choices on the temporal discounting task (Koloski, Hulyalkar, et al., 2024) (electrode locations, **Fig 2A**, **Table 1**). At the time of reward outcome (0-1s after reward onset) we found populations of electrodes that were significantly activated or suppressed for reward (small and large reward at 0.5s delays) (**Fig 2B**, **Table 1**). Reward-related activations in beta power were strongest in orbitofrontal and striatal regions, whereas thalamic and hippocampal regions showed suppression (**Fig 2B**, **Table 1**). Only activated electrodes were analyzed further. The group of suppressed electrodes (reminiscent of a default-mode-like network) likely represents a distinct network that deserves to be explored further.

### Activity on Small Reward Trials

We hypothesized that methylphenidate may exert influence over beta reward signals to modulate cost-benefit decision-making. The shift in behavior to prefer delayed reward following methylphenidate could be achieved by 1) decreasing value of immediate reward or 2) increasing value of large reward, despite temporal delays. For the reward-activated electrodes, we first examined their activity on small reward trials. High-dose methylphenidate significantly reduced beta power during small reward trials (main effect of drug; F_(3,581.01)_ =2.90*, p=.034*) (**Fig 3A, B**) – an effect observed at all electrode locations (**Fig 3C**). Beta suppression may reduce the salience of immediate rewards, biasing animals toward waiting.

**Figure 3:**
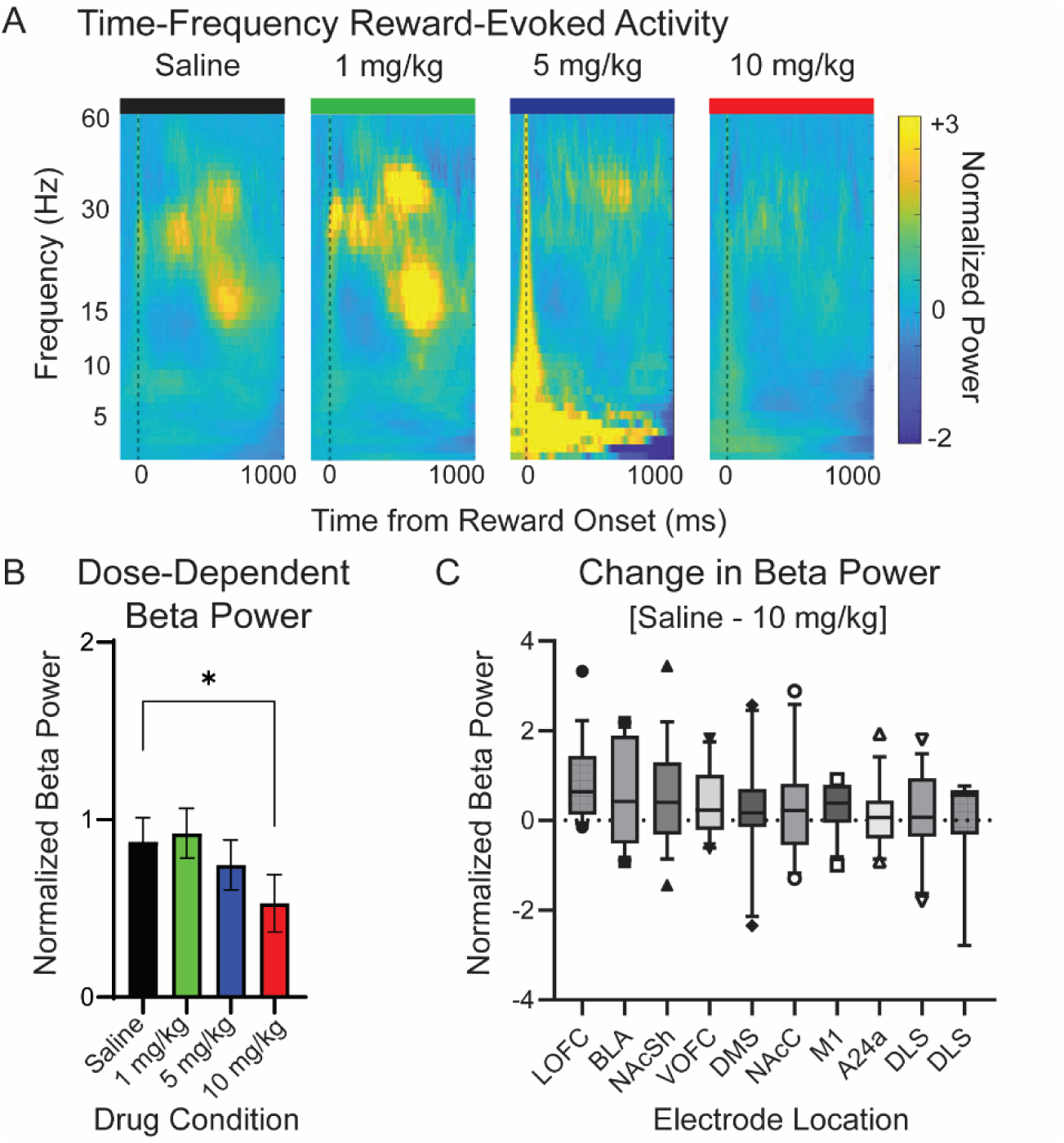
Reward-evoked activity on small reward trials. A) Time-frequency plots of reward-evoked LFP activity in lateral orbitofrontal cortex (LOFC) under different drug conditions. Power is normalized to baseline and time-locked to reward onset (dashed line). Methylphenidate (5 and 10 mg/kg) suppressed high-frequency activity (beta) during small rewards. B) Dose-dependent effects of methylphenidate on reward-evoked beta power (averaged across 10 electrodes; main effect of drug; F_(3,581.01)_ =2.90*, p=.034*). Compared with saline, beta power was significantly reduced following 10mg/kg methylphenidate (t_(316)_=2.21, *p=.028*). C) Box-and-whisker plots (10-90%) showing electrode-specific differences in beta suppression from saline to 10mg/kg. While methylphenidate reduced beta activity on all electrodes, the magnitude of suppression was greater in LOFC, basolateral amygdala, and nucleus accumbens (main effect of electrode; F_(9,615.50)_ =2.90*, p=.034*). Bar graphs show Mean± SEM. Significance is denoted with asterisks (* *p*<.05, ** *p*<.01; \*\*\**p*<.001). Abbreviations are listed in **Table 1**.

### Activity on Large Reward Trials

Reward-evoked beta activity was measured on the same electrodes during large reward trials under delay conditions (0.5, 2, 5, 10, 20s). Under saline conditions, beta power declined with increasing delay, replicating prior findings (main effect of delay; F_(4,2667.20)_ =20.47*, p<.001*) (**Fig 4A**). Methylphenidate attenuated this decline, with high doses restoring reward-evoked beta activity even at 10-20s delays (main effect of dose; F_(3, 2666.41)_ =7.72*, p<.001*) (**Fig. 4 B-D**). At the 5s delay, where drug effects on choice were strongest, the difference in power between small and large rewards was minimized (**Fig 4E**). Methylphenidate simultaneously suppresses beta power for small reward while restoring beta power for large, delayed reward, consistent with the shift in behavior biasing waiting for reward.

**Figure 4:**
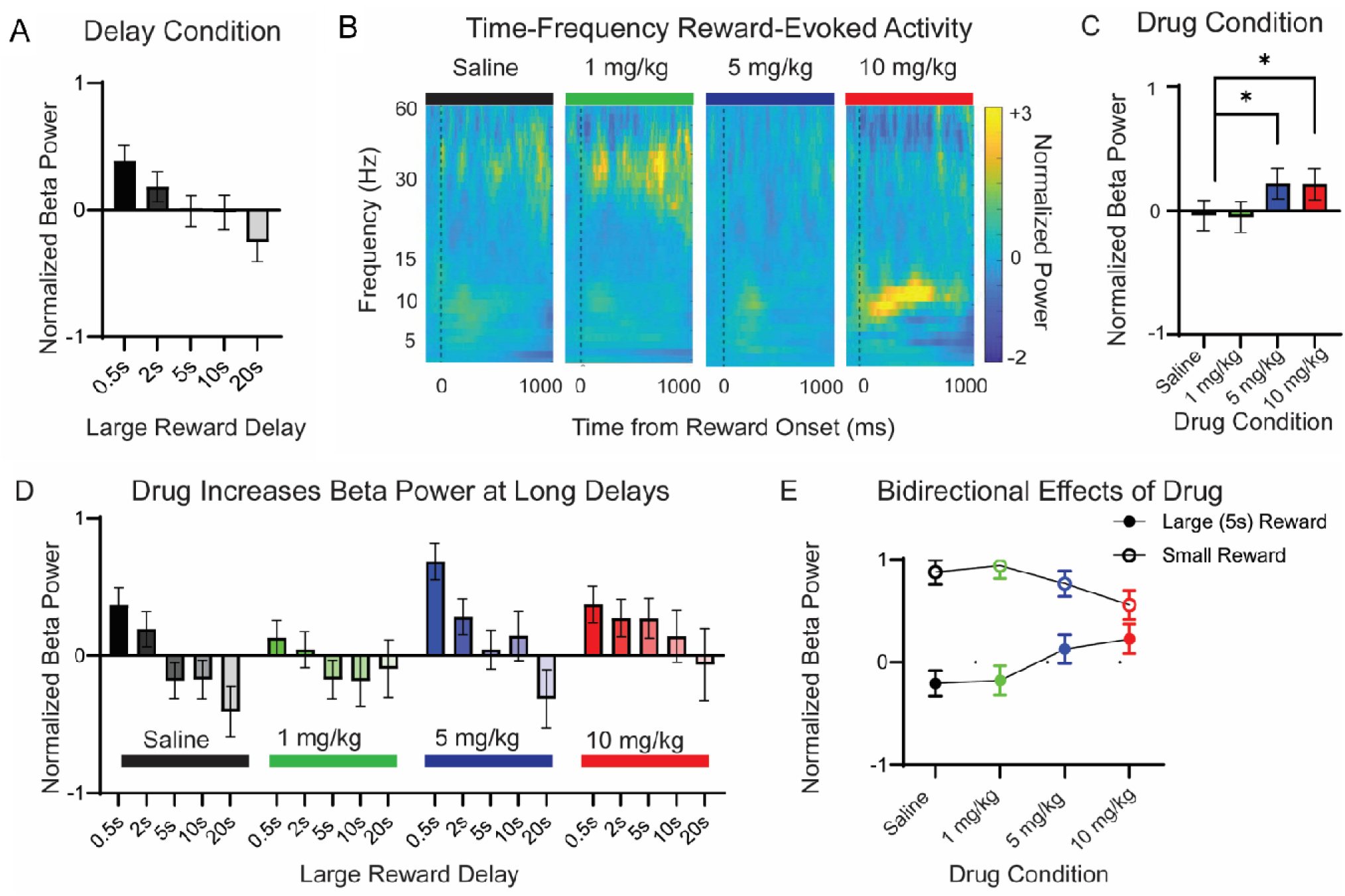
Reward-evoked activity on large reward trials. A) Beta power is delay-dependent (main effect of delay; F_(4,2667.20)_ =20.47*, p<.001*), with greater suppression at longer reward delays. B) Time-frequency plots of reward-evoked LOFC under different drug conditions. Power is normalized to baseline and time-locked to reward onset (dashed line). Under saline conditions, large reward delivery suppressed evoked-beta power, whereas methylphenidate (10mg/kg) increased beta oscillations. C) Dose-dependent effects of methylphenidate on reward-evoked beta power (averaged across 10 electrodes; main effect of dose; F_(3, 2666.41)_ =7.72*, p<.001*). Compared with saline, both the 5 and 10 mg/kg doses significantly increased beta power (saline v. 5mg/kg, t_(1821)_= -3.94, *p=.038*; saline v. 10 mg/kg, t_(1641)_=-3.05, *p=.041*). D) Beta power across temporal delays (0.5, 2, 5, 10, 20s) for each drug condition (saline, 1, 5, and 10mg/kg). A significant dose x delay interaction was observed F_(12, 2663.23)_ =2.32*, p=.006*). Post hoc tests showed that with 10mg/kg methylphenidate, beta power no longer differed across delays (F_(4,472.30)_=1.279, *p=.277*). Large doses of methylphenidate reduce delay-related beta suppression. E) At the 5s delay condition (the temporal delay with the strongest behavior effect; see **Fig 1C**), we compared the effects of methylphenidate on small and large reward trials. Methylphenidate had a dual effect: increasing beta power for the large, delayed reward while suppressing beta power for the small, immediate reward. All plots show Mean ± SEM. Box and whisker plots depict the 10-90% range. Significance is denoted with asterisks (* *p*<.05, ** *p*<.01; \*\*\**p*<.001).

### Neurobehavioral Correlations and Network Activity

Finally, we examined how methylphenidate influenced network connectivity by measuring weighted phase lag index (wPLI) between LOFC and the other reward-activated regions (10 connectivity pairs). LOFC showed distinct connectivity patterns with the other regions during small and large (5s delay) rewards that shifted dynamically with methylphenidate. LOFC connectivity (strongest with BLA and vOFC) during small reward trials, was reduced by methylphenidate (main effect of drug F_(3,640)_=4.09, *p=.007*) (**Fig 5A**), potentially weakening immediate-reward drive. During large reward trials, LOFC connectivity was less affected (F_(3,444)_=0.877, *p=0.66*) (**Fig 5B**). There was a trend for LOFC-BLA connectivity to dose-dependently increase and LOFC-VMS connectivity to decrease (perhaps minimizing the overall effect of drug, due to their opposition).

**Figure 5:**
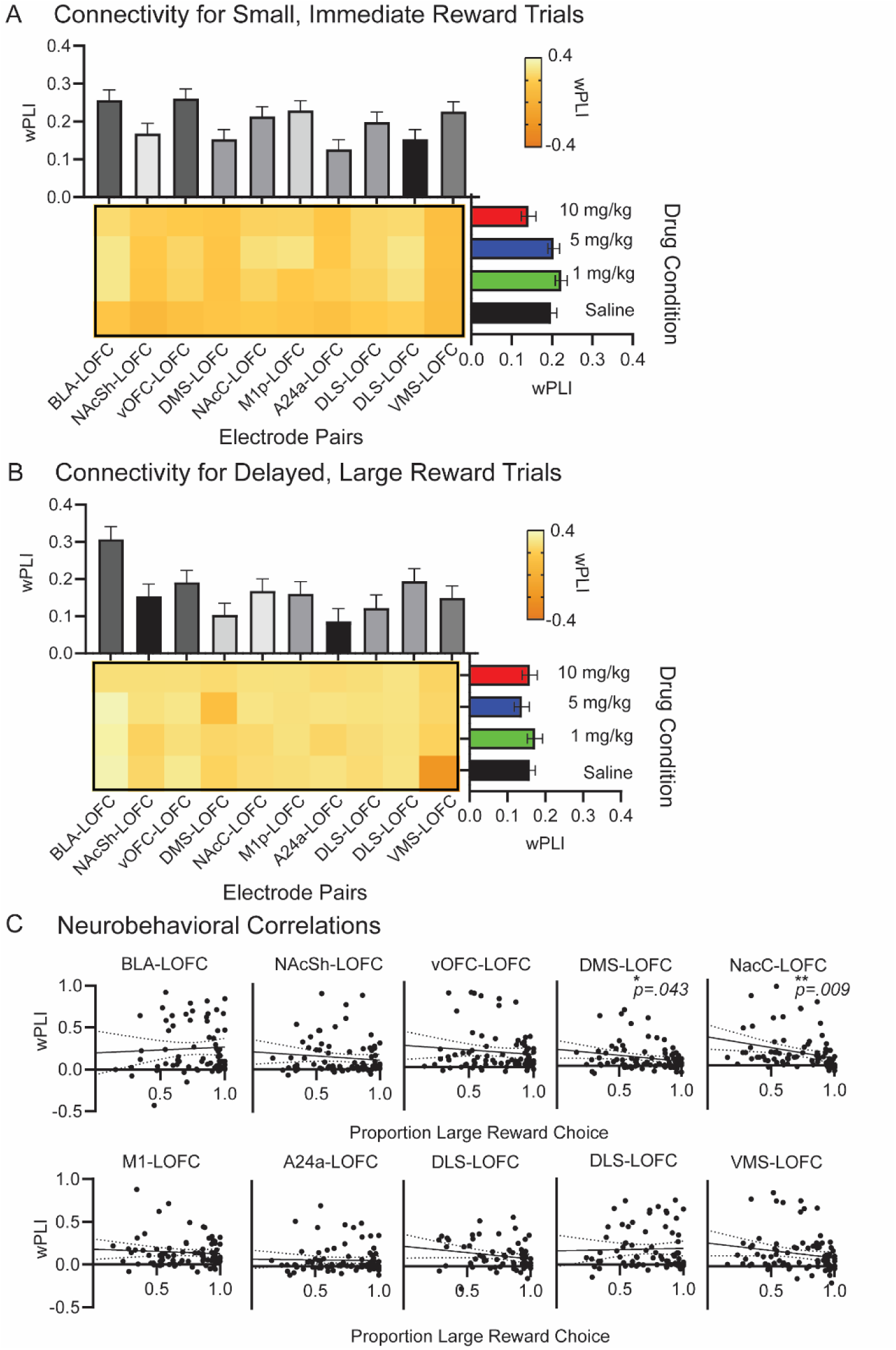
Neurobehavioral correlations with cortico-striatal connectivity. A) Connectivity patterns between lateral orbitofrontal cortex (LOFC) and other reward- activated regions across drug conditions. Heatmaps show beta frequency weighted phase lag index (wPLI) for each LOFC-electrode pair under saline and drug conditions (wPLI range from -.4 (dark, weak connectivity) to +.4 (light, strong connectivity)). Bar plots summarize Mean ± SEM wPLI for each electrode pair (columns) and each drug condition (rows). 10mg/kg methylphenidate reduced connectivity overall (main effect of drug F_(3,640)_=4.09, *p=.007*). B) Connectivity patterns during delayed, large reward trials. No statistically significant differences were observed (F_(3,444)_=0.877, *p=0.66*), but trends included increased LOFC-BLA connectivity and decreased LOFC-VMS connectivity with methylphenidate. C) Neurobehavioral correlations between connectivity and choice behavior. For each LOFC-electrode pair (10 pairs), the relationship between wPLI during small reward trials and the proportion of large reward choices at the 5s delay was assessed using Pearson’s correlation fit to a linear regression. Significant negative correlations were observed for DMS-LOFC (R^2^ = 4.22, *p=.043*) and NacC-LOFC (R^2^ = 7.10, *p=.009*) electrode pairs., indicating that higher cortico-striatal connectivity during small reward choice predicted fewer large reward choices.

To explore the relationship between network activity and behavior, we looked for correlations between connectivity on small reward trials and the proportion of large reward choices (5s delay) for each session, regardless of drug condition. Although weak, most regions had an inverse relationship between small reward connectivity and large reward preference (DMS (R^2^ = 4.22, *p=.043*) and NacC (R^2^ = 7.10, *p=.009*) had significant negative correlations). Reduced connectivity for small rewards correlated with greater preference for delayed rewards (**Fig 5C**). Methylphenidate’s ability to reduce impulsivity on this task, and promote waiting, is associated with less cortico- striatal connectivity and reward-evoked power for small rewards.

## Discussion

Methylphenidate is well known to reduce impulsive choice in temporal discounting task across clinical and preclinical models (Castrellon et al., 2021; Daood et al., 2022; Floresco, 2013; Martinez et al., 2020), but treatment response is highly variable (Caprioli et al., 2015; Clatworthy et al., 2009; Coghill & Banaschewski, 2009; Daood et al., 2022; Rajala et al., 2015; Westbrook et al., 2020). Identifying a neurophysiological marker associated with impulsivity that predicts treatment response would therefore have significant clinical utility. Here, we demonstrate that cortico-striatal beta oscillations could provide such a marker. On a delay discounting task, methylphenidate reduced impulsivity by decreasing beta power and network connectivity for small, immediate rewards, while simultaneously enhancing beta power for large, delayed rewards. These shifts in oscillatory dynamics were associated with an increased tendency to choose the larger, but delayed, reward option. Our findings support the idea that reward-evoked beta oscillations may serve as a translational biomarker of treatment responsivity to monoaminergic drugs such as methylphenidate.

Across species, beta oscillations in prefrontal and striatal regions track reward expectancy, subjective value, and choice selection (Cohen et al., 2007; HajiHosseini & Holroyd, 2015; Hoy et al., 2023; Koloski, Hulyalkar, et al., 2024, 2024; Marco-Pallarés et al., 2015; Patai et al., 2022; Xiao et al., 2024; Zavala et al., 2015). Our data extends this literature by demonstrating that methylphenidate modulates reward-evoked beta oscillations in a direction consistent with reduced impulsivity: away from immediate rewards and towards larger, delayed rewards (**Fig. 3**, **Fig. 4**). Reduced connectivity for small rewards was inversely correlated with delayed reward choice, suggesting that oscillatory activity within cortico-striatal circuits directly influences discounting behavior **(Fig. 5**).

Prefrontal cortex and its ventral striatum targets are central to temporal discounting (De Water et al., 2017; Winstanley et al., 2005). Inactivation of striatal PFC projections increases impulsive choice (Wenzel et al., 2023), and patients with impulse disorders present altered cortico-striatal functional connectivity (Hong et al., 2015). Our findings align with models of cost-benefit decision-making, in which immediate rewards engage mesolimbic valuation circuits, while delayed rewards engage frontoparietal control regions (McClure et al., 2004; Varma et al., 2023). Prior studies similarly report that cortical beta power is elevated during non-impulsive choices, whereas theta oscillations are linked with impulsive responding (Gui et al., 2019; Stam et al., 2024). In comparison to cortical beta, theta oscillations increase following negative feedback (Marco-Pallarés et al., 2015; Phaf & Rotteveel, 2012) signal effort requirements (Seamans et al., 2024), or anticipate reward (Knudsen & Wallis, 2020; Marquardt et al., 2017). Interestingly, even sensorimotor beta oscillations have been implicated in motivational drive (Chen & Kwak, 2022; Verhein et al., 2023), suggesting that methylphenidate’s effects may extend beyond valuation circuits.

It was not clear how methylphenidate would alter brain signals associated with impulsive choice. We find that methylphenidate had a bidirectional influence on cortico- striatal network activity dampening impulsive signals for small, immediate rewards while enhancing signals large, delayed reward. These findings align with human fMRI studies showing that methylphenidate enhances frontrostriatal connectivity during delayed, non-impulsive choices, the magnitude of which predicts behavioral improvement in patients (Daood et al., 2022).. Methylphenidate also reduces intra-thalamic/ cortico-thalamic coupling (Gorka et al., 2020), consistent with a mechanism that “frees” prefrontal valuation networks from competing thalamic drive, reducing stimulus-driven responses and prioritizing goal-directed valuation. In addition to within-band effects, methylphenidate may also influence cross-frequency interactions. In children with ADHD, greater theta/beta and delta/beta power ratios predict symptom improvement (Sari Gokten et al., 2019), and in adults, methylphenidate decreases low/high frequency coupling (Zammit & Muscat, 2023), suggesting that the balance between slower and faster oscillatory rhythms is another key mechanism by which drug shapes network activity and impulsive behavior.

Methylphenidate increases extracellular dopamine and norepinephrine, and our findings suggest that beta oscillations may serve as a network-level signature of this monoaminergic modulation. From a dopaminergic perspective, beta oscillations may index tonic rather than phasic signaling. Phasic dopamine signals encode reward prediction errors (RPE)- transient increases in striatal activity that occur when outcomes exceed expectation (Schultz, 2016). Large phasic responses in ventral striatum are linked to impulsive responding when reward expectancy is low (Cools, 2006). By elevating tonic dopamine, methylphenidate dampens phasic responses, decreasing RPEs and reducing impulsive choice (Evers et al., 2017; Li et al., 2024). Our observation that beta activity is high when reward expectancy is high and low when it is low (Koloski, Hulyalkar, et al., 2024) is consistent with the idea that beta reflects tonic dopaminergic tone.

Norepinephrine also likely contributes to this framework. Methylphenidate promotes cortical “up states’ and reduces slow oscillations, effects also observed with selective norepinephrine reuptake inhibitors (Shen & Shi, 2021). Given the sparse dopaminergic innervation of prefrontal cortex, dopamine released from norepinephrine terminals may play a critical role in regulating cortical excitability (Arnsten, 2009; Morón et al., 2002). Thus, our findings likely reflect joint dopamine and norepinephrine modulation of cortico- striatal circuits.

Opposed to directly influencing cost-benefit decision-making, another pathway by which methylphenidate may reduce impulsivity is through altered time perception. In humans, methylphenidate leads to interval underestimation, which could make delayed rewards subjectively feel closer in time (and thus more valuable) (Farias et al., 2019). Time interval underestimation was associated with reduced alpha power in attention networks (Farias et al., 2019), consistent with reports that methylphenidate decreases insular activation and spatial attention bias (Ivanov et al., 2014; Peled-Avron et al., 2024).

While our study focused on the reward outcome period, future work should analyze oscillatory dynamic during the delay or choice period to better understand whether altered temporal processing/ attention contributes to reduced impulsivity.

Our findings provide a mechanistic link between monoaminergic modulation, reward valuation networks, and behavior. Methylphenidate reduces impulsive choice by modulating cortico-striatal beta oscillations- dampening immediate-reward signals and enhancing delayed-reward signals. Given the heterogeneity of methylphenidate efficacy in ADHD and related impulsivity disorders (Cortese et al., 2018), finding beta oscillations as a marker of drug effects has high translational potential. EEG measures, such as beta/theta power ratios, already can predict clinical response to methylphenidate in children with ADHD (Arns et al., 2018; Sari Gokten et al., 2019). Extending this framework, beta oscillations could be developed as a circuit-level assay to identify responders, guide dosing, and evaluate new therapeutics for impulsivity-related disorders.

## References

Ainslie, G. W. (1974). IMPULSE CONTROL IN PIGEONS^1^. Journal of the Experimental Analysis of Behavior, 21(3), 485–489. 10.1901/jeab.1974.21-485

Arns, M., Vollebregt, M. A., Palmer, D., Spooner, C., Gordon, E., Kohn, M., Clarke, S., Elliott, G. R., & Buitelaar, J. K. (2018). Electroencephalographic biomarkers as predictors of methylphenidate response in attention-deficit/hyperactivity disorder. European Neuropsychopharmacology, 28(8), 881–891. 10.1016/j.euroneuro.2018.06.002

Arnsten, A. F. T. (2009). Stress signalling pathways that impair prefrontal cortex structure and function. Nature Reviews Neuroscience, 10(6), 410–422. 10.1038/nrn2648

Askenasy, E. P., Taber, K. H., Yang, P. B., & Dafny, N. (2007). METHYLPHENIDATE (RITALIN): BEHAVIORAL STUDIES IN THE RAT. International Journal of Neuroscience, 117(6), 757–794. 10.1080/00207450600910176

Baarendse, P. J. J., & Vanderschuren, L. J. M. J. (2012). Dissociable effects of monoamine reuptake inhibitors on distinct forms of impulsive behavior in rats. Psychopharmacology, 219(2), 313–326. 10.1007/s00213-011-2576-x

Basar, K., Sesia, T., Groenewegen, H., Steinbusch, H. W. M., Visser-Vandewalle, V., & Temel, Y. (2010). Nucleus accumbens and impulsivity. Progress in Neurobiology, 92(4), 533–557. 10.1016/j.pneurobio.2010.08.007

Buscher, N., Ojeda, A., Francoeur, M., Hulyalkar, S., Claros, C., Tang, T., Terry, A., Gupta, A., Fakhraei, L., & Ramanathan, D. S. (2020). Open-source raspberry Pi-based operant box for translational behavioral testing in rodents. Journal of Neuroscience Methods, 342, 108761. 10.1016/j.jneumeth.2020.108761

Caprioli, D., Jupp, B., Hong, Y. T., Sawiak, S. J., Ferrari, V., Wharton, L., Williamson, D. J., McNabb, C., Berry, D., Aigbirhio, F. I., Robbins, T. W., Fryer, T. D., & Dalley, J. W. (2015). Dissociable Rate-Dependent Effects of Oral Methylphenidate on Impulsivity and D_2/3_ Receptor Availability in the Striatum. The Journal of Neuroscience, 35(9), 3747–3755. 10.1523/JNEUROSCI.3890-14.2015

Cardinal, R. N., Parkinson, J. A., Hall, J., & Everitt, B. J. (2002). Emotion and motivation: The role of the amygdala, ventral striatum, and prefrontal cortex. Neuroscience & Biobehavioral Reviews, 26(3), 321–352. 10.1016/S0149-7634(02)00007-6

Castrellon, J. J., Meade, J., Greenwald, L., Hurst, K., & Samanez-Larkin, G. R. (2021). Dopaminergic modulation of reward discounting in healthy rats: A systematic review and meta-analysis. Psychopharmacology, 238(3), 711–723. 10.1007/s00213-020-05723-5

Chen, X.-J., & Kwak, Y. (2022). Contribution of the sensorimotor beta oscillations and the cortico-basal ganglia-thalamic circuitry during value-based decision making: A simultaneous EEG-fMRI investigation. NeuroImage, 257, 119300. 10.1016/j.neuroimage.2022.119300

Chudasama, Y. (2011). Animal models of prefrontal-executive function. Behavioral Neuroscience, 125(3), 327–343. 10.1037/a0023766

Clatworthy, P. L., Lewis, S. J. G., Brichard, L., Hong, Y. T., Izquierdo, D., Clark, L., Cools, R., Aigbirhio, F. I., Baron, J.-C., Fryer, T. D., & Robbins, T. W. (2009). Dopamine Release in Dissociable Striatal Subregions Predicts the Different Effects of Oral Methylphenidate on Reversal Learning and Spatial Working Memory. The Journal of Neuroscience, 29(15), 4690–4696. 10.1523/JNEUROSCI.3266-08.2009

Coghill, D., & Banaschewski, T. (2009). The genetics of attention-deficit/hyperactivity disorder. Expert Review of Neurotherapeutics, 9(10), 1547–1565. 10.1586/ern.09.78

Cohen, M. X., Elger, C. E., & Ranganath, C. (2007). Reward expectation modulates feedback-related negativity and EEG spectra. NeuroImage, 35(2), 968–978. 10.1016/j.neuroimage.2006.11.056

Cools, R. (2006). Dopaminergic modulation of cognitive function-implications for l-DOPA treatment in Parkinson’s disease. Neuroscience & Biobehavioral Reviews, 30(1), 1–23. 10.1016/j.neubiorev.2005.03.024

Cortese, S., Adamo, N., Del Giovane, C., Mohr-Jensen, C., Hayes, A. J., Carucci, S., Atkinson, L. Z., Tessari, L., Banaschewski, T., Coghill, D., Hollis, C., Simonoff, E., Zuddas, A., Barbui, C., Purgato, M., Steinhausen, H.-C., Shokraneh, F., Xia, J., & Cipriani, A. (2018). Comparative efficacy and tolerability of medications for attention-deficit hyperactivity disorder in children, adolescents, and adults: A systematic review and network meta-analysis. The Lancet Psychiatry, 5(9), 727– 738. 10.1016/S2215-0366(18)30269-4

Dalley, J. W., Everitt, B. J., & Robbins, T. W. (2011). Impulsivity, Compulsivity, and Top-Down Cognitive Control. Neuron, 69(4), 680–694. 10.1016/j.neuron.2011.01.020

Daood, M., Peled-Avron, L., Ben-Hayun, R., Nevat, M., Aharon-Peretz, J., Tomer, R., & Admon, R. (2022). Fronto-striatal connectivity patterns account for the impact of methylphenidate on choice impulsivity among healthy adults. Neuropharmacology, 216, 109190. 10.1016/j.neuropharm.2022.109190

De Water, E., Mies, G. W., Figner, B., Yoncheva, Y., Van Den Bos, W., Castellanos, F. X., Cillessen, A. H. N., & Scheres, A. (2017). Neural mechanisms of individual differences in temporal discounting of monetary and primary rewards in adolescents. NeuroImage, 153, 198–210. 10.1016/j.neuroimage.2017.04.013

De Wit, H., Wade, T. R., & Richards, J. B. (2000). Effects of dopaminergic drugs on delayed reward as a measure of impulsive behavior in rats. Psychopharmacology, 150(1), 90–101. 10.1007/s002130000402

Evenden, J. L., & Ryan, C. N. (1996). The pharmacology of impulsive behaviour in rats: The effects of drugs on response choice with varying delays of reinforcement. Psychopharmacology, 128(2), 161–170. 10.1007/s002130050121

Evers, E. A., Stiers, P., & Ramaekers, J. G. (2017). High reward expectancy during methylphenidate depresses the dopaminergic response to gain and loss. Social Cognitive and Affective Neuroscience, 12(2), 311–318. 10.1093/scan/nsw124

Faraone, S. V. (2018). The pharmacology of amphetamine and methylphenidate: Relevance to the neurobiology of attention-deficit/hyperactivity disorder and other psychiatric comorbidities. Neuroscience and Biobehavioral Reviews, 87, 255–270. 10.1016/j.neubiorev.2018.02.001

Farias, T. L., Marinho, V., Carvalho, V., Rocha, K., Da Silva, P. R. A., Silva, F., Teles, A. S., Gupta, D., Ribeiro, P., Velasques, B., Cagy, M., Bastos, V. H., Silva-Junior, F., & Teixeira, S. (2019). Methylphenidate modifies activity in the prefrontal and parietal cortex accelerating the time judgment. Neurological Sciences, 40(4), 829–837. 10.1007/s10072-018-3699-1

Floresco, S. B. (2013). Prefrontal dopamine and behavioral flexibility: Shifting from an “inverted-U” toward a family of functions. Frontiers in Neuroscience, 7. 10.3389/fnins.2013.00062

Floresco, S. B., Tse, M. T. L., & Ghods-Sharifi, S. (2008). Dopaminergic and Glutamatergic Regulation of Effort- and Delay-Based Decision Making. Neuropsychopharmacology, 33(8), 1966–1979. 10.1038/sj.npp.1301565

Francoeur, M. J., Tang, T., Fakhraei, L., Wu, X., Hulyalkar, S., Cramer, J., Buscher, N., & Ramanathan, D. R. (2021). Chronic, Multi-Site Recordings Supported by Two Low-Cost, Stationary Probe Designs Optimized to Capture Either Single Unit or Local Field Potential Activity in Behaving Rats. Frontiers in Psychiatry, 12, 678103. 10.3389/fpsyt.2021.678103

Gorka, A. X., Lago, T. R., Balderston, N., Torrisi, S., Fuchs, B., Grillon, C., & Ernst, M. (2020). Intrinsic connections between thalamic sub-regions and the lateral prefrontal cortex are differentially impacted by acute methylphenidate. Psychopharmacology, 237(6), 1873–1883. 10.1007/s00213-020-05505-z

Gui, D.-Y., Yu, T., Hu, Z., Yan, J., & Li, X. (2019). Author Correction: Dissociable functional activities of cortical theta and beta oscillations in the lateral prefrontal cortex during intertemporal choice. Scientific Reports, 9(1), 12141. 10.1038/s41598-019-48220-2

Haber, S. N., & Knutson, B. (2010). The Reward Circuit: Linking Primate Anatomy and Human Imaging. Neuropsychopharmacology, 35(1), 4–26. 10.1038/npp.2009.129

HajiHosseini, A., & Holroyd, C. B. (2015). Sensitivity of frontal beta oscillations to reward valence but not probability. Neuroscience Letters, 602, 99–103. 10.1016/j.neulet.2015.06.054

Hamilton, K. R., Mitchell, M. R., Wing, V. C., Balodis, I. M., Bickel, W. K., Fillmore, M., Lane, S. D., Lejuez, C. W., Littlefield, A. K., Luijten, M., Mathias, C. W., Mitchell, S. H., Napier, T. C., Reynolds, B., Schütz, C. G., Setlow, B., Sher, K. J., Swann, A. C., Tedford, S. E., … Moeller, F. G. (2015). Choice impulsivity: Definitions, measurement issues, and clinical implications. *Personality Disorders: Theory*, Research, and Treatment, 6(2), 182–198. 10.1037/per0000099

Hong, S.-B., Harrison, B. J., Fornito, A., Sohn, C.-H., Song, I.-C., & Kim, J.-W. (2015). Functional dysconnectivity of corticostriatal circuitry and differential response to methylphenidate in youth with attention-deficit/hyperactivity disorder. Journal of Psychiatry and Neuroscience, 40(1), 46–57. 10.1503/jpn.130290

Hoy, C. W., De Hemptinne, C., Wang, S. S., Harmer, C. J., Apps, M. A. J., Husain, M., Starr, P. A., & Little, S. (2023). Beta and theta oscillations track effort and previous reward in human basal ganglia and prefrontal cortex during decision making. 10.1101/2023.12.05.570285

Ivanov, I., Liu, X., Clerkin, S., Schulz, K., Fan, J., Friston, K., London, E. D., Schwartz, J., & Newcorn, J. H. (2014). Methylphenidate and brain activity in a reward/conflict paradigm: Role of the insula in task performance. European Neuropsychopharmacology, 24(6), 897–906. 10.1016/j.euroneuro.2014.01.017

Kable, J. W., & Glimcher, P. W. (2007). The neural correlates of subjective value during intertemporal choice. Nature Neuroscience, 10(12), 1625–1633. 10.1038/nn2007

Knudsen, E. B., & Wallis, J. D. (2020). Closed-Loop Theta Stimulation in the Orbitofrontal Cortex Prevents Reward-Based Learning. Neuron, 106(3), 537–547.e4. 10.1016/j.neuron.2020.02.003

Koloski, M. F., Hulyalkar, S., Barnes, S. A., Mishra, J., & Ramanathan, D. S. (2024). Cortico-striatal beta oscillations as a reward-related signal. *Cognitive, Affective*, & Behavioral Neuroscience, 24(5), 839–859. 10.3758/s13415-024-01208-6

Koloski, M. F., Terry, A., Lee, N., & Ramanathan, D. S. (2024). Methylphenidate, but not citalopram, decreases impulsive choice in rats performing a temporal discounting task. Frontiers in Psychiatry, 15, 1385502. 10.3389/fpsyt.2024.1385502

Li, Y., Huang, Y., Chen, J. J., Hyland, B. I., & Wickens, J. R. (2024). Phasic dopamine signals are reduced in the spontaneously hypertensive rat and increased by methylphenidate. European Journal of Neuroscience, 59(7), 1567–1584. 10.1111/ejn.16269

MacKillop, J., Amlung, M. T., Few, L. R., Ray, L. A., Sweet, L. H., & Munafò, M. R. (2011). Delayed reward discounting and addictive behavior: A meta-analysis. Psychopharmacology, 216(3), 305–321. 10.1007/s00213-011-2229-0

Marco-Pallarés, J., Münte, T. F., & Rodríguez-Fornells, A. (2015). The role of high-frequency oscillatory activity in reward processing and learning. Neuroscience & Biobehavioral Reviews, 49, 1–7. 10.1016/j.neubiorev.2014.11.014

Marquardt, K., Sigdel, R., & Brigman, J. L. (2017). Touch-screen visual reversal learning is mediated by value encoding and signal propagation in the orbitofrontal cortex. Neurobiology of Learning and Memory, 139, 179–188. 10.1016/j.nlm.2017.01.006

Martinez, E., Pasquereau, B., Drui, G., Saga, Y., Météreau, É., & Tremblay, L. (2020). Ventral striatum supports Methylphenidate therapeutic effects on impulsive choices expressed in temporal discounting task. Scientific Reports, 10(1), 716. 10.1038/s41598-020-57595-6

McClure, S. M., Laibson, D. I., Loewenstein, G., & Cohen, J. D. (2004). Separate Neural Systems Value Immediate and Delayed Monetary Rewards. Science, 306(5695), 503–507. 10.1126/science.1100907

Mitchell, S. H. (2019). Linking Delay Discounting and Substance Use Disorders: Genotypes and Phenotypes. Perspectives on Behavior Science, 42(3), 419–432. 10.1007/s40614-019-00218-x

Morón, J. A., Brockington, A., Wise, R. A., Rocha, B. A., & Hope, B. T. (2002). Dopamine Uptake through the Norepinephrine Transporter in Brain Regions with Low Levels of the Dopamine Transporter: Evidence from Knock-Out Mouse Lines. The Journal of Neuroscience, 22(2), 389–395. 10.1523/JNEUROSCI.22-02-00389.2002

Patai, E. Z., Foltynie, T., Limousin, P., Akram, H., Zrinzo, L., Bogacz, R., & Litvak, V. (2022). Conflict Detection in a Sequential Decision Task Is Associated with Increased Cortico-Subthalamic Coherence and Prolonged Subthalamic Oscillatory Response in the β Band. The Journal of Neuroscience, 42(23), 4681– 4692. 10.1523/JNEUROSCI.0572-21.2022

Pearson, D. A., Santos, C. W., Casat, C. D., Lane, D. M., Jerger, S. W., Roache, J. D., Loveland, K. A., Lachar, D., Faria, L. P., Payne, C. D., & Cleveland, L. A. (2004). Treatment Effects of Methylphenidate on Cognitive Functioning in Children With Mental Retardation and ADHD. Journal of the American Academy of Child & Adolescent Psychiatry, 43(6), 677–685. 10.1097/01.chi.0000124461.81324.13

Peled-Avron, L., Daood, M., Ben-Hayun, R., Nevat, M., Aharon-Peretz, J., Admon, R., & Tomer, R. (2024). Methylphenidate reduces spatial attentional bias by modulating fronto-striatal connectivity. Cerebral Cortex, 34(9), bhae379. 10.1093/cercor/bhae379

Perry, W., Minassian, A., Paulus, M. P., Young, J. W., Kincaid, M. J., Ferguson, E. J., Henry, B. L., Zhuang, X., Masten, V. L., Sharp, R. F., & Geyer, M. A. (2009). A Reverse-Translational Study of Dysfunctional Exploration in Psychiatric Disorders: From Mice to Men. Archives of General Psychiatry, 66(10), Article 10. 10.1001/archgenpsychiatry.2009.58

Peters, S. K., Dunlop, K., & Downar, J. (2016). Cortico-Striatal-Thalamic Loop Circuits of the Salience Network: A Central Pathway in Psychiatric Disease and Treatment. Frontiers in Systems Neuroscience, 10. 10.3389/fnsys.2016.00104

Phaf, R. H., & Rotteveel, M. (2012). Affective Monitoring: A Generic Mechanism for Affect Elicitation. Frontiers in Psychology, 3. 10.3389/fpsyg.2012.00047

Pujara, M., & Koenigs, M. (2014). Mechanisms of Reward Circuit Dysfunction in Psychiatric Illness: Prefrontal–Striatal Interactions. The Neuroscientist, 20(1), 82–95. 10.1177/1073858413499407

Rajala, A. Z., Jenison, R. L., & Populin, L. C. (2015). Decision making: Effects of methylphenidate on temporal discounting in nonhuman primates. Journal of Neurophysiology, 114(1), 70–79. 10.1152/jn.00278.2015

Rodriguez, M. L., & Logue, A. W. (1988). Adjusting delay to reinforcement: Comparing choice in pigeons and humans. Journal of Experimental Psychology. Animal Behavior Processes, 14(1), 105–117.

Roesch, M. R., & Bryden, D. W. (2011). Impact of Size and Delay on Neural Activity in the Rat Limbic Corticostriatal System. Frontiers in Neuroscience, 5. 10.3389/fnins.2011.00130

Roffman, J. L., & Raskin, L. A. (1997). Stereotyped Behavior: Effects of d-Amphetamine and Methylphenidate in the Young Rat. Pharmacology Biochemistry and Behavior, 58(4), 1095–1102. 10.1016/S0091-3057(97)00321-3

Sari Gokten, E., Tulay, E. E., Beser, B., Elagoz Yuksel, M., Arikan, K., Tarhan, N., & Metin, B. (2019). Predictive Value of Slow and Fast EEG Oscillations for Methylphenidate Response in ADHD. Clinical EEG and Neuroscience, 50(5), 332–338. 10.1177/1550059419863206

Schultz, W. (2016). Dopamine reward prediction error coding. Dialogues in Clinical Neuroscience, 18(1), 23–32. 10.31887/DCNS.2016.18.1/wschultz

Seamans, J. K., Emberly, E., White, S., Morningstar, M., Linsenbardt, D., Ma, B., Czachowski, C. L., & Lapish, C. C. (2024). Neural basis of cognitive control signals in anterior cingulate cortex during delay discounting. Cold Spring Harbor Laboratory. 10.1101/2024.06.07.597894

Shen, G., & Shi, W. (2021). Amphetamine promotes cortical Up state: Role of adrenergic receptors. Addiction Biology, 26(1). 10.1111/adb.12879

Slezak, J. M., & Anderson, K. G. (2011). Effects of acute and chronic methylphenidate on delay discounting. Pharmacology Biochemistry and Behavior, 99(4), 545–551. 10.1016/j.pbb.2011.05.027

Stam, C. H., Van Der Veen, F. M., & Franken, I. H. A. (2024). Evidence for post-decisional conflict monitoring in delay discounting. Biological Psychology, 192, 108849. 10.1016/j.biopsycho.2024.108849

Torrecillos, F., Tinkhauser, G., Fischer, P., Green, A. L., Aziz, T. Z., Foltynie, T., Limousin, P., Zrinzo, L., Ashkan, K., Brown, P., & Tan, H. (2018). Modulation of Beta Bursts in the Subthalamic Nucleus Predicts Motor Performance. The Journal of Neuroscience, 38(41), 8905–8917. 10.1523/JNEUROSCI.1314-18.2018

Van Den Bos, W., & McClure, S. M. (2013). TOWARDS A GENERAL MODEL OF TEMPORAL DISCOUNTING. Journal of the Experimental Analysis of Behavior, 99(1), 58–73. 10.1002/jeab.6

Van Den Bos, W., Rodriguez, C. A., Schweitzer, J. B., & McClure, S. M. (2014). Connectivity Strength of Dissociable Striatal Tracts Predict Individual Differences in Temporal Discounting. Journal of Neuroscience, 34(31), 10298–10310. 10.1523/JNEUROSCI.4105-13.2014

Van Den Bosch, R., Lambregts, B., Määttä, J., Hofmans, L., Papadopetraki, D., Westbrook, A., Verkes, R.-J., Booij, J., & Cools, R. (2022). Striatal dopamine dissociates methylphenidate effects on value-based versus surprise-based reversal learning. Nature Communications, 13(1), 4962. 10.1038/s41467-022-32679-1

Varma, M. M., Zhen, S., & Yu, R. (2023). Not all discounts are created equal: Regional activity and brain networks in temporal and effort discounting. NeuroImage, 280, 120363. 10.1016/j.neuroimage.2023.120363

Verhein, J. R., Vyas, S., & Shenoy, K. V. (2023). Methylphenidate modulates motor cortical dynamics and behavior. 10.1101/2023.10.15.562405

Volkow, N. D. (1997). Effects of methylphenidate on regional brain glucose metabolism in humans: Relationship to dopamine D2 receptors. American Journal of Psychiatry, 154(1), 50–55. 10.1176/ajp.154.1.50

Wenzel, J. M., Zlebnik, N. E., Patton, M. H., Smethells, J. R., Ayvazian, V. M., Dantrassy, H. M., Zhang, L.-Y., Mathur, B. N., & Cheer, J. F. (2023). Selective chemogenetic inactivation of corticoaccumbal projections disrupts trait choice impulsivity. Neuropsychopharmacology, 48(12), 1821–1831. 10.1038/s41386-023-01604-5

Westbrook, A., Van Den Bosch, R., Määttä, J. I., Hofmans, L., Papadopetraki, D., Cools, R., & Frank, M. J. (2020). Dopamine promotes cognitive effort by biasing the benefits versus costs of cognitive work. Science, 367(6484), 1362–1366. 10.1126/science.aaz5891

Winstanley, C. A., Eagle, D. M., & Robbins, T. W. (2006). Behavioral models of impulsivity in relation to ADHD: Translation between clinical and preclinical studies. Clinical Psychology Review, 26(4), 379–395. 10.1016/j.cpr.2006.01.001

Winstanley, C. A., Theobald, D. E. H., Dalley, J. W., & Robbins, T. W. (2005). Interactions between Serotonin and Dopamine in the Control of Impulsive Choice in Rats: Therapeutic Implications for Impulse Control Disorders. Neuropsychopharmacology, 30(4), 669–682. 10.1038/sj.npp.1300610

Xiao, J., Adkinson, J. A., Myers, J., Allawala, A. B., Mathura, R. K., Pirtle, V., Najera, R., Provenza, N. R., Bartoli, E., Watrous, A. J., Oswalt, D., Gadot, R., Anand, A., Shofty, B., Mathew, S. J., Goodman, W. K., Pouratian, N., Pitkow, X., Bijanki, K. R., … Sheth, S. A. (2024). Beta activity in human anterior cingulate cortex mediates reward biases. Nature Communications, 15(1), 5528. 10.1038/s41467-024-49600-7

Yang, P. B., Swann, A. C., & Dafny, N. (2006). Dose-response characteristics of methylphenidate on locomotor behavior and on sensory evoked potentials recorded from the VTA, NAc, and PFC in freely behaving rats. 2(1), 3. 10.1186/1744-9081-2-3

Zammit, N., & Muscat, R. (2023). Alpha/beta-gamma decoupling in methylphenidate medicated ADHD patients. Frontiers in Neuroscience, 17, 1267901. 10.3389/fnins.2023.1267901

Zavala, B., Damera, S., Dong, J. W., Lungu, C., Brown, P., & Zaghloul, K. A. (2015). Human Subthalamic Nucleus Theta and Beta Oscillations Entrain Neuronal Firing During Sensorimotor Conflict. *Cerebral Cortex*, bhv244. 10.1093/cercor/bhv244

